# Estimating the duration of perceptual moments from early infancy to adulthood

**DOI:** 10.64898/2026.10.02.756257

**Authors:** Sara Dochez, Emilie Serraille, Liuba Papeo, Flavia Mengarelli, Marie-Maude Geoffray-Cassar, Jean-Rémy Hochmann

**Affiliations:** CNRS UMR5229 – Institut des Sciences Cognitives Marc Jeannerod, 67 Bd Pinel 69675 Bron, France; Université Claude Bernard Lyon 1, France; CH Le Vinatier – Bron, France; RESHAPE, RESearch on HealthcAre PErformance, INSERM U1290, France

## Abstract

Although human perception appears continuous, converging evidence suggests that sensory input is integrated over discrete temporal windows. The duration of these Temporal Integration Windows (TIWs) determines the temporal resolution of perception, but how this fundamental property develops across infancy and childhood remains unknown. Here, we estimate TIW duration from 5 months of age to adulthood using a novel, instruction-free eye-tracking paradigm that isolates pre-attentive visual integration. Across four experiments with the same paradigm (N = 308 participants), we measured spontaneous detection of a visual target that emerges only when two alternating displays are integrated within a single perceptual window. We observed a striking developmental shortening of the TIW. In adults and 5-year-olds, integration succeeded at short interstimulus intervals (ISIs; 16–32 ms) but failed at longer intervals (83-116 ms), yielding an estimated TIW of ∼57 ms. At 3 years, the TIW extended to ∼100 ms. In infancy, integration remained robust at substantially longer ISIs, with estimates of ∼250 ms at 18 and 8 months, and ∼400 ms at 5 months. The steepest reduction occurred within the first year of life. These findings reveal an approximately sevenfold acceleration in the temporal resolution of visual perception from early infancy to adulthood. Progressive shortening of TIWs suggests that infants and adults integrate sensory information over fundamentally different temporal scales, with implications for perceptual organization, learning, and neurodevelopmental trajectories.

## INTRODUCTION

Our experience of the visual world appears continuous; but this is an illusion. Perception consists of a sequence of *moments* ^1–3^ that result from a discrete sampling of the continuous stream of sensory information ^4^. The speed of this sampling controls the temporal resolution of perception, which in turn constitutes a fundamental organizing principle of our representation of the world, constraining the encoding of temporal order and the detection of causal and social interactions ^5,6^.

How long does a moment last? Presumably, its duration relates to the speed of information processing in our brains, and the speed of communication between neurons. In the course of development, progressive myelination accelerates these communications, predicting an acceleration of information processing, and a reduction of a moment duration. In the current study, we operationalize a moment as a time window in which successive incoming visual information is combined into a single coherent event: the *Temporal Integration Window (TIW)*. In contrast, two events occurring in separate TIWs are perceived as two segmented events. While previous empirical studies have yielded variable TIWs in infants ^7,8^, children ^9,10^ and adults ^9,11^, none have pursued a systematic developmental investigation using the same paradigm across the full lifespan, from early infancy to adulthood. Here, we employ the same paradigm with populations from 5 months of age to adulthood, in order to quantify the variation of the TIW duration in the course of development.

To enable investigations in young preverbal infants, we designed a paradigm that does not require any verbal instruction, feedback to participants, nor any explicit responses. We manipulated the speed of alternation between two displays that can generate a unique attractive stimulus (i.e., one full face among an array of half-faces), when the two displays are shown close enough in time to be integrated into a single percept. The longest temporal interval at which displays form this unique percept is exactly the TIW. Gaze position, as measured with an eye-tracking system, was used to evaluate the duration of the TIW at various ages. In Experiment 1, we showed that our paradigm replicates previous estimations of the TIW in adults and demonstrated that the TIW we are probing relates to the pre-attentional stage of perception. In Experiments 2-4, we evaluated the duration of the TIW in infants and children from 5 months to 5 years of age, highlighting important and systematic variations in the course of development.

## RESULTS

### Experiment 1

In our paradigm, stimuli consisted in the alternation of two displays (Display A and Display B; Figure 1). Each display showed a number of grayscales faces or half-faces arranged at equidistance from the center of the screen, on a black background. In *integration trials*, each display contained only half-faces. Only one of the half-faces of Display B was positioned to complement one of the half-faces of Display A, so that if the two displays were integrated into one unique percept, participants should see one full face (hereafter the target) among half-faces (Figure 1.a). In *segmentation trials*, Display A contained half-faces at all non-target (distractor) locations and no item at the target location. Display B contained the complementary half-faces at the non-target locations and one full face at the target location. If the two displays were integrated, participants should perceive full faces at all locations (Figure 1b). If the displays were segmented, they should perceive the single full face in Display B. In both integration and segmentation trials, the perception of a full face among half-faces should capture attention, eliciting longer looks towards the target than towards the distractors.

**Figure 1.**
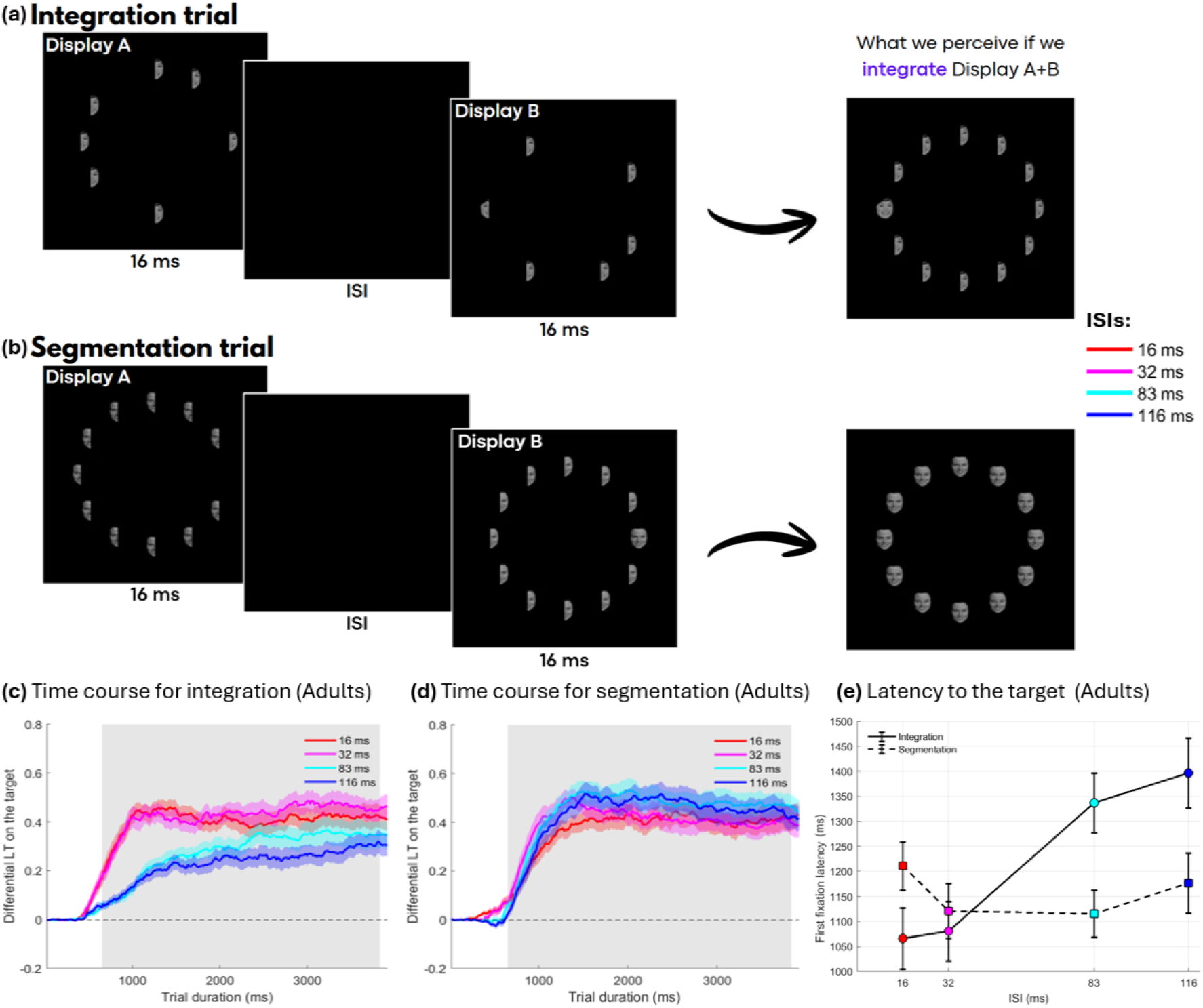
Paradigm and results of Experiments 1. **(a)** In integration trials, Displays A and B alternate, each containing half-faces. One pair of complementary half-faces forms a full face when the displays are integrated. **(b)** In segmentation trials, Display A contains half-faces at the non-target locations and no item at the target location, whereas Display B contains the complementary half-faces and one full face at the target location. Integrating the displays produces full faces at all locations, whereas segmenting them reveals the single full face in Display B. **(c–e)** Results of Experiment 1 in adults (N = 26). **(c–d)** Time course of target detection in integration **(c)** and segmentation **(d)** trials at ISIs of 16, 32, 83, and 116 ms. Target detection was assessed by comparing gaze toward the target location with gaze toward the distractor locations. Grey shading indicates the Trial Type × ISI interaction window (650–3910 ms). **(e)** Mean first-fixation latency to the target as a function of trial type and ISI.

**Figure 2.**
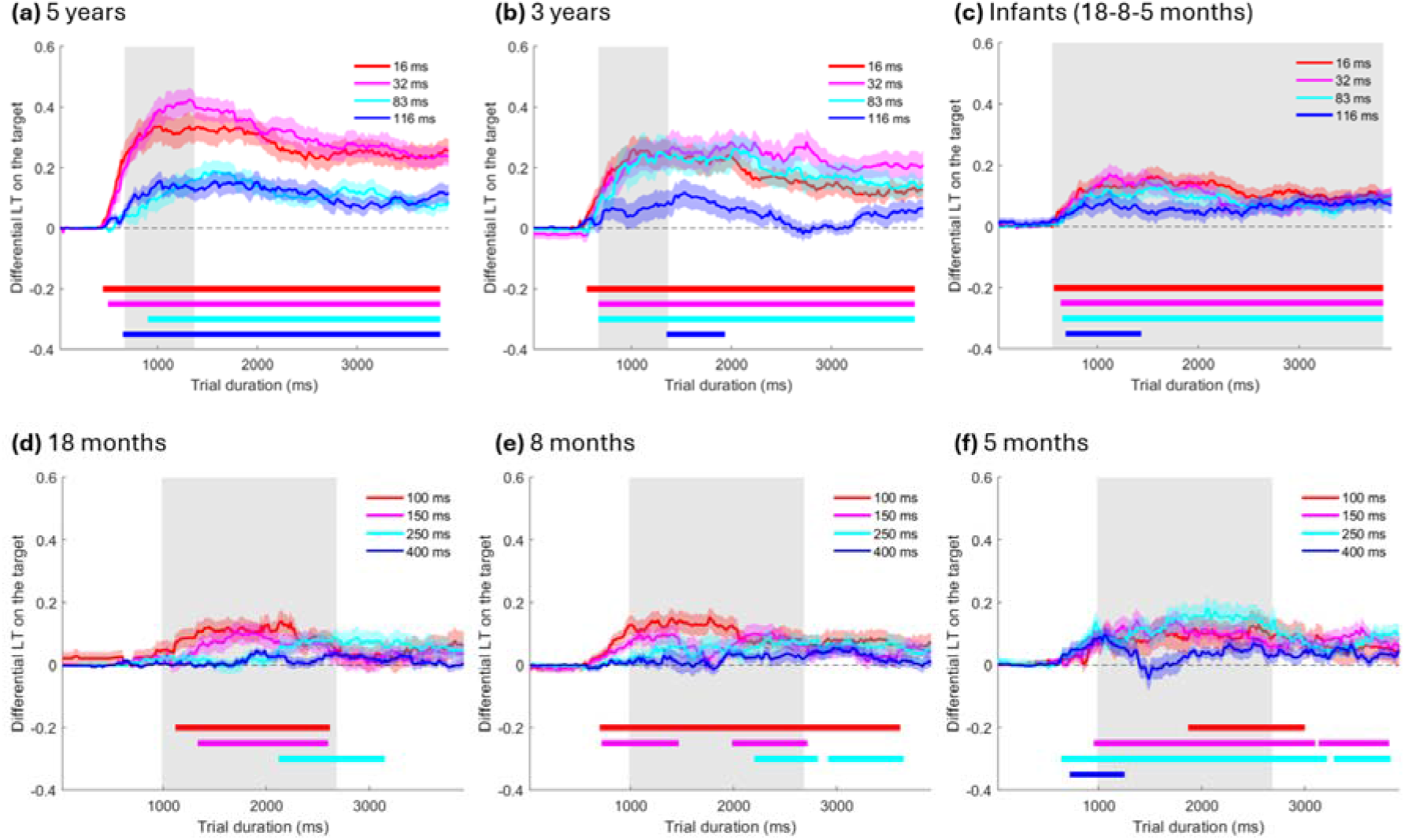
Differential Looking Times time course for integration trials in Experiments 2-4. **(a-c)** Results of Experiments 2 and 3. Target detection was assessed by comparing gaze toward the target location with gaze toward the opposite location. **(a)** Time course of target detection in integration trials for 5-year-olds (N = 26) at ISIs of 16, 32, 83, and 116 ms. **(b)** Time course for 3-year-olds (N = 26) at the same ISIs. **(c)** Time course for 5-, 8-, and 18-month-olds (total N = 81) at the same ISIs. In **(a)** and **(b)**, grey shading indicates the Age × Trial Type × ISI interaction window (666–1216 ms). In **(c)**, it indicates the interval showing a main effect of Trial Type. **(d-f)** Results of Experiment 4. **(d)** Time course of target detection in integration trials for 18-month-olds (N = 48), **(e)** 8-month-olds (N = 50), and **(f)** 5-month-olds (N = 51), at ISIs of 100, 150, 250, and 400 ms. In **(a-f)**, colored horizontal bars indicate significant time intervals for each ISI (cluster-mass permutation tests, p < 0.05). In **(d-f)**, grey shading indicates the window showing main effects of Trial Type and ISI (983–2683 ms).

Note that, in integration trials, the target could theoretically be detected in absence of integration, because the target half-face in Display B had a different left–right orientation from the distractor half-faces. This is however unlikely due to perception constancy in front of symmetrical transformations. Moreover, the inverted target should be more salient as alternation slows down, thereby predicting better target detection at longer ISIs; i.e., the opposite pattern expected from temporal integration. In segmentation trials, the target could be detected even in presence of integration on the basis of a different flicker pattern: the target location alternated between a full face and an empty location, whereas the distractor locations alternated between left and right half-faces. This difference should be detectable even at the fastest tested alternation rates (60 Hz) ^12^, and therefore does not predict an increase of target detection at slower alternation rates.

In experience 1, we manipulated the number of distractors; Displays showed either 6 items (1 target full-face among 5 half-faces) or 12 items (1 target full-face among 11 half-faces). The target full-face could appear at any position. Each trial began when participants fixated a central stimulus, and lasted 4 seconds. The crucial variable was the interstimulus interval (ISI) between Display A and Display B: 16, 32, 83, or 116 ms.

Twenty-six adults participated in Experiment 1 (16 females; mean age = 27.16 years, SD = 5.68) and were tested on 20 trials per condition. In the following analyses, we only considered trials where participants looked at the screen for at least 50% of the first 2 seconds of the trial. Participants contributed on average 19.01 (SD= 0.09) trials per condition.

First, we tested whether the target detection in our paradigm lies in the preattentional stage of perception. According to the Feature Integration Theory ^13^, visual perception unfolds in two sequential stages: an early, pre-attentive stage in which basic visual features are processed automatically and in parallel across the visual field, and a later, attention-dependent stage in which features are bound into coherent object representations. In the pre-attentive stage, a target differing from distractors along a basic feature dimension automatically captures attention, a phenomenon known as pop-out, independently of the number of distractors. This set-size independence distinguishes pre-attentive pop-out from the serial, attention-demanding search that characterizes the later stage. We thus analyzed whether participants first looked at the target or at a distractor, how fast they found the target, and whether this varied with the number of distractors. Note that there is a long debate whether faces per se pop-out ^14–17^. In the present design, the target could pop-out on the basis of low-level information, especially size or roundedness, but crucially, only if Displays A and B are integrated in integration trials, and only if Displays A and B are segmented in segmentation trials.

Across all experimental conditions (integration and segmentation trials, all ISIs), participants looked at the target first in 38.9% of trials. To test if this exceeded chance, we computed for each participant the proportion of first fixations directed to the target, separately for trials with 5 or 11 distractors. In 5 distractor trials, participants fixated the target first in 36.6 % ± 2.6 % (SEM) of trials, significantly above the theoretical chance level of 16.7 % *(t(25) = 6.76, p < .0001)*. In 11 distractor trials, the proportion of first fixations to the target reached 41.7 % ± 2.2 %, also significantly above the theoretical chance level of 8.3 % *(t(25) = 14.58, p < .0001)*.

We then examined whether the number of distractors influenced the latency to fixate the target. Mean latency on the target was 1328 ms (SD = 243 ms) in the 5-distractor condition, and 1309 ms (SD = 182 ms) in the 11-distractor condition. A paired-samples t-test revealed no significant difference between conditions, *t(25) = 0.68, p = 0.505*, Cohen’s d = 0.13, BF01=3.92. These results suggest that the number of distractors did not affect how rapidly participants oriented toward the target, providing evidence that the targets captured visual attention and hence that the integration process we are studying occurs in the pre-attentive stage of perception.

Having established that the target detection lies in the early preattentional stage of perception, we evaluated the Temporal Integration Window by testing the effects of ISIs (16, 32, 83, 116 ms) and Trial Type on participants’ target detection.

#### Latencies

We analyzed the latency to fixate the target. A repeated-measures ANOVA with Trial Type and ISIs as within-subject factors revealed a significant main effect of Trial Type (F(1,25) = 5.21, *p = .0313*), with longer latencies in integration compared to segmentation trials. There was also a significant main effect of ISIs (F(3,75) = 7.40, *p < .001*). Most importantly, the interaction Trial Type x ISIs was significant (F(3,75) = 9.08, *p < .001*). In integration trials we found shorter latencies at 32 compared to 83 ms t(25) = −3.50, p = 0.002, no significant differences were noted between 16 and 32 or between 83 and 116 ms (all *p > 0.42*). No significant difference was found for segmentation trials.

#### Time course of gaze fixation

For each participant, at each time point, we computed the Differential Looking Time (DLT), defined as: *DLT(t)= (Proportion of trials looking at the target at time t) – (Mean Proportion of trials looking at any distractor at time t)*. We conducted cluster-based permutation tests on the time-course of DLT, relying on a repeated-measures ANOVA with Trial Type (integration, segmentation) and ISI (16, 32, 83, 116 ms) as within-subject factors (Figures 1.c and 1.d). This analysis revealed a significant main effect of Trial Type between 900 ms and 3150 ms after stimulus presentation *(p = 0.003),* and a main effect of ISIs between 333 ms and 1460 ms (*p = 0.013*). Most importantly, the Trial Type × ISIs interaction was also significant, from 650 ms to 3910 ms *(p < 0.001)*, confirming that the impact of ISIs on target detection differed between integration and segmentation trials.

Post-hoc tests restricted to this interaction window (650–3910 ms) revealed no significant differences for ISIs of 16 ms and 32 ms *(t (25)= −1.28, p = 0.2109)*, but a significant decline in DLT between ISI of 32 ms and ISI of 83 ms *(t(25) = 5.94, p < 0.001)*; and between ISI of 83 ms and ISI of 116 ms (*t(25)= 2.38, p = 0.0253)*. The reverse effect was observed for segmentation performance with no significant differences between ISIs of 16 ms and 32 ms *(t (25)= −1.83, p = 0.079)*, a significant increase in DLT between ISIs of 32 and 83 ms (t(25) = −2.98, p= 0.006); and no differences for ISIs of 83 ms and 116 ms *(t (25)= 0.98, p = 0.3381)*. Overall, these analyses converge in showing that, in integration trials, the detection of the full-face target gets easier with shorter ISI, especially when moving from 83 to 32 ms. In parallel, it gets more difficult in segmentation trials. We can thus estimate that the TIW in adults closes between 32 and 83 ms, that is approximately around 57 ms, congruent with prior estimates with a similar paradigm ^9,11^.

### EXPERIMENT 2

The stimuli and procedure of Experiment 2 were identical to those of Experiment 1, except that the number of distractors was maintained at 11 and the target could only appear in two positions, symmetrically located on the central axis of the screen.

Fifty-two typically developing children, divided in one group of 5-year-olds (N=26, 14 females, mean age= 64.94 months, SD=3.4 months), and one group of 3-year-olds (N=26, 12 females, mean age= 41.78 months, SD=3.69 months), provided on average 10.64 (SD= 0.29) trials per condition (Table S1).

As evidenced in Tables S2-S3, the number of trials in which children looked at the target at least once varied substantially across conditions, compromising the validity of the latency analysis. In Experiments 2-4 with children and infants, we thus focused on the analysis of Differential Looking Times (DLTs).

As for adults in Experiment 1, we computed the time course of the Differential Looking Time (DLT) for each ISI, for integration and for segmentation trials. Cluster mass permutation tests relying on repeated-measures ANOVA assessed the effects of Age (3, 5 years) as between-subjects factor, and Trial Type (integration, segmentation) and ISIs (16, 32, 83, 116 ms) as within-subject factors on the propensity to gaze at the target (DLTs). A significant main effect of Trial Type was detected between 583 ms and 3916 ms (p <0.001); a significant main effect of ISIs was detected between 533 ms and 3916 ms (*p <0.001*) and a significant main effect of Age was detected between 566 ms and 3916 ms (*p <0.001*). In addition, a significant Trial Type × ISIs interaction was observed between 900 and 3916 ms (*p < 0.001*). No significant interactions were found between Age and ISIs or between Age and Trial Type. Most importantly, a significant triple interaction between Age, ISIs and Trial Type was observed between 666 ms to 1216 ms (*p = .027*).

To explain the triple interaction, we averaged the difference between the integration and segmentation DLTs in the identified time window (666–1216 ms) for each ISI condition and each participant. For each age group, we assessed the effect of ISIs with a repeated-measures ANOVA. An effect of ISIs was observed for 5-year-olds (*F*(3,75) = 11.31, *p* < .0001). Pairwise comparisons indicated a marked decline between ISIs of 32 and 83 ms (*t(25) = 4.88, p < .001*), whereas no reliable difference emerged between ISIs of 16 and 32 ms (*t(25) = −1.45, p = 0.159*) or between ISIs of 83 and 116 ms (*t(25) = −1.86, p = 0.075*). The effect of ISI was not significant for 3-year-olds (*F*(3,75) = 2.23, *p* = .104). Pairwise comparisons showed no differences between ISIs of 16 and 32 ms (t(25) = 0.49, p = 0.626) or between ISIs of 32 and 83 ms (t(25) = −1.04, p = 0.307). A significant difference was present between ISIs of 83 and 116 ms (*t(25)=2.47, p= 0.0209*).

#### Target detection in each condition

We further ran planned cluster-mass permutation tests on each ISI and each Trial Type separately to determine whether participants successfully detected the target. Results are summarized in Table 1. Five-year-olds detected the target for all ISIs for both integration trials and segmentation trials. Three-year-olds, in contrast, detected the target for all ISIs in integration trials, though the detection appeared weaker and later for ISI of 116 ms. In segmentation trials, 3-year-olds only detected the target for ISI of 83 and 116 ms, with earlier detection for the longer ISI.

**Table 1.** Evidence for target detection in Experiment 2.

| Age group | ISI | Significant clusters in integration trials | Segmentation trials |
| --- | --- | --- | --- |
| 5 years | 16 ms | 450–3916 ms ($p < 0.001$) | 633–3733 ms ($p < 0.001$) |
| | 32 ms | 533–3916 ms ($p < 0.001$) | 1116–1833 ms ($p = 0.002$);<br>2950–3516 ms ($p = 0.0291$) |
| | 83 ms | 583–3916 ms ($p < 0.001$) | 650–4000 ms ($p < 0.001$) |
| | 116 ms | 500–3916 ms ($p < 0.001$) | 783–3500 ms ($p = 0.001$) |
| 3 years | 16 ms | 550–4000 ms ($p < 0.001$) | No significant clusters |
| | 32 ms | 666–4000 ms ($p < 0.001$) | No significant clusters |
| | 83 ms | 666–4000 ms ($p < 0.001$) | 1300–4000 ms ($p < 0.001$) |
| | 116 ms | 1350–1933 ms ($p = 0.047$) | 900–2083 ms ($p = 0.006$) |

Overall, combining all analyses, we can estimate the TIW to last about 57 ms at 5 years (i.e., between 32 and 83 ms), similar to that of adults. At 3 years, the TIW appears longer, as evidenced by the good detection of the target with ISI of 83 ms in integration trials. Target detection appears weaker for ISI of 116 ms, suggesting a TIW closing only around 100 ms (i.e., between 83 and 116 ms).

### EXPERIMENT 3

The stimuli and procedure of Experiment 3 were identical to those of Experiment 2, except for the age of participants. Eighty-one typically developing infants of 18 months (N=23, 14 females, mean age= 18.15 months, SD=0.28 months), 8 months (N=27, 10 females, mean age= 8.35 months, SD=0.23 months), or 5 months (N=31, 11 females, mean age= 5.25 months, SD=0.29 months) provided on average 5.58 (SD= 0.28) trials per condition (Table S1).

As in Experiments 1-2, we computed the time course of Differential Looking Time (DLT) for each ISI, for integration and for segmentation trials. Cluster mass permutation tests relying on repeated-measures ANOVA assessed the effects of Age (5, 8, 18 months) as between-subjects factor, and Trial Type (integration, segmentation) and ISIs (16, 32, 83, 116 ms) as within-subject factors, on the propensity to gaze at the target (DLTs). These analyses revealed a main effect of Trial Type, with a large significant cluster extending from 550 to 3916 ms after stimulus onset (p < .001), indicating stronger target detection in integration than in segmentation trials. No significant effects of ISIs or Age were observed, and no interaction between factors.

#### Target detection in each condition

We further ran planned cluster-mass permutation tests on each condition separately to determine whether participants successfully detected the target. Results are summarized in Table 2. Infants, 5- to 18-months of age, detected the target for all ISIs in integration trials, but never in segmentation trials. Overall, these analyses suggest that infants aged 5 to 18 months integrate visual information across ISIs up to at least 116 ms, suggesting a TIW substantially longer than that of adults and pre-school children. To gain better estimate of the TIW in young infants, Experiment 4 assessed infant’s detection of the target for ISIs varying between 100 and 400 ms.

**Table 2.**
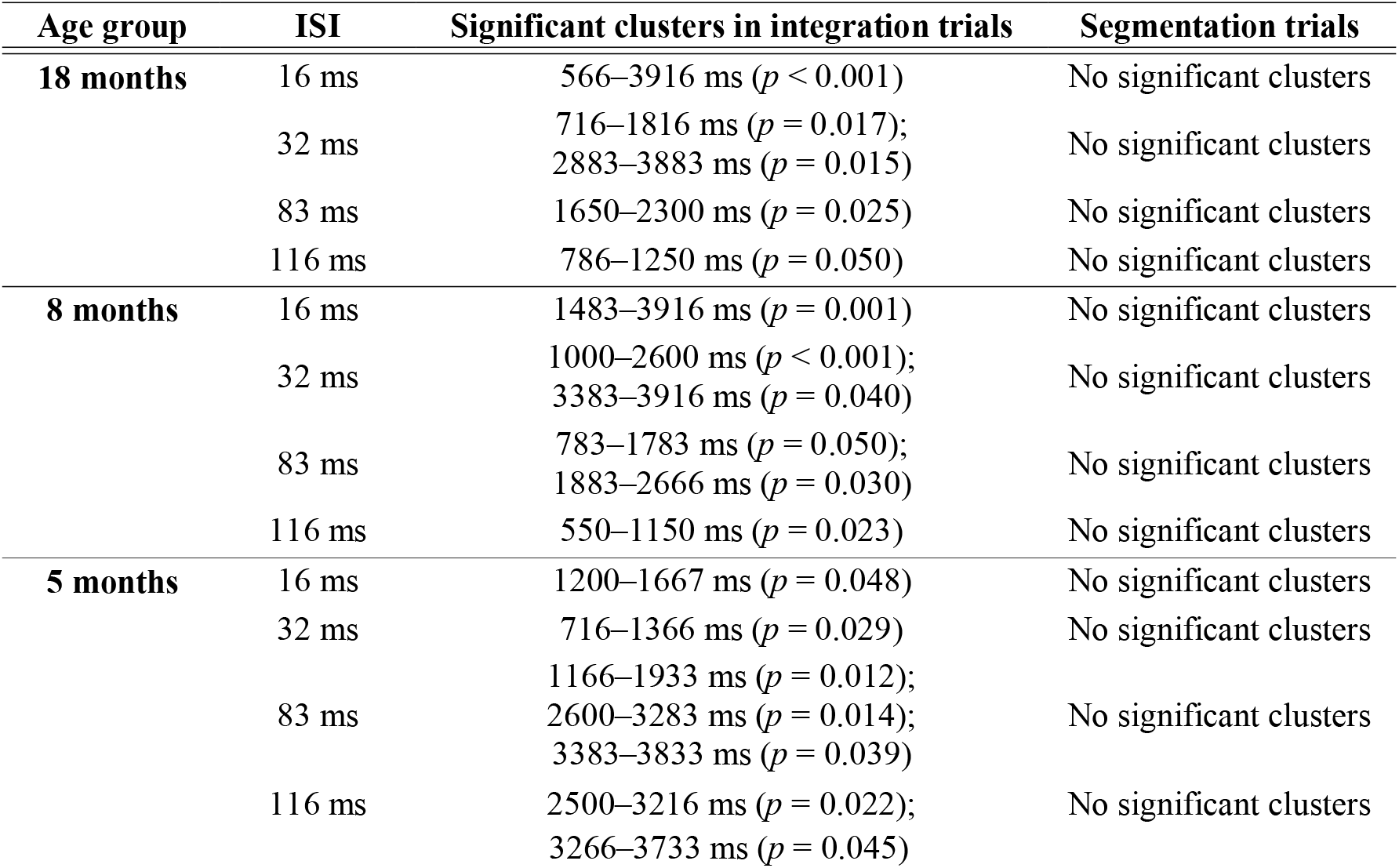
Evidence for target detection in Experiment 3.

### EXPERIMENT 4

Stimuli and procedure were identical to those of Experiments 2 and 3, except that longer interstimulus intervals (ISIs: 100, 150, 250, and 400 ms) were used. One-hundred-forty-nine typically developing infants of 18 months (N=48, 27 females, mean age= 18.6 months, SD= 0.31 months), 8 months (N=50, 25 females, mean age= 8.5 months, SD=0.32 months), or 5 months (N=51, 19 females, mean age= 5.3 months, SD=0.30 months) contributed an average of 6.28 (SD= 0.32) trials per condition (Table S1).

As in Experiments 1-3, we computed the time course of Differential Looking Time (DLT) for each ISI, for integration and for segmentation trials. Cluster-mass permutation tests relying on repeated-measures ANOVA assessed the effects of Age (5, 8, 18 months) as between-subjects factor, Trial Type (integration, segmentation), and ISI (100, 150, 250, 400 ms) as within-subject factors, on the propensity to gaze at the target (DLTs). A main effect of Trial Type was observed from 717 to 3916 ms (p < .001), reflecting stronger target detection in integration than in segmentation trials across all infant groups. A main effect of ISI was observed from 983 to 2683 ms (p = .0002). No significant main effect of Age nor any significant interaction between factors was found.

#### Target detection in each condition

We further ran cluster-mass permutation tests on each condition separately to determine whether participants successfully detected the target. Results are summarized in Table 3. They confirm that the TIW in infancy substantially exceeds 100 ms across all tested ages (5 to 18 months), and further suggest a developmental gradient within the first year of life. At 8 and 18 months, we observe target detection in integration trials for ISIs of 100,150 and 250 ms, but not for ISIs of 400 ms. The detection appears relatively late for ISI of 250 ms, suggesting a weaker pop-out effect. Overall this suggests the TIW may be closing around 250 ms at these ages. In contrast, 5-month-olds detected the target in integration trials for all ISIs 100, 150, 250 and 400 ms, suggesting a TIW at 5 months of at least 400 ms. Supporting these conclusions, cluster mass permutation tests showed no significant effect of ISI on target detection in either integration or segmentation trials at 5 months. At 8 months, a significant effect of ISI was observed in integration trials between 817 and 1283 ms (*p* = 0.0126), whereas no significant cluster was found in segmentation trials. The same pattern was observed at 18 months, with a significant ISI effect in integration trials between 1283 and 2017 ms (*p* = 0.0106), but no significant cluster in segmentation trials.

**Table 3.**
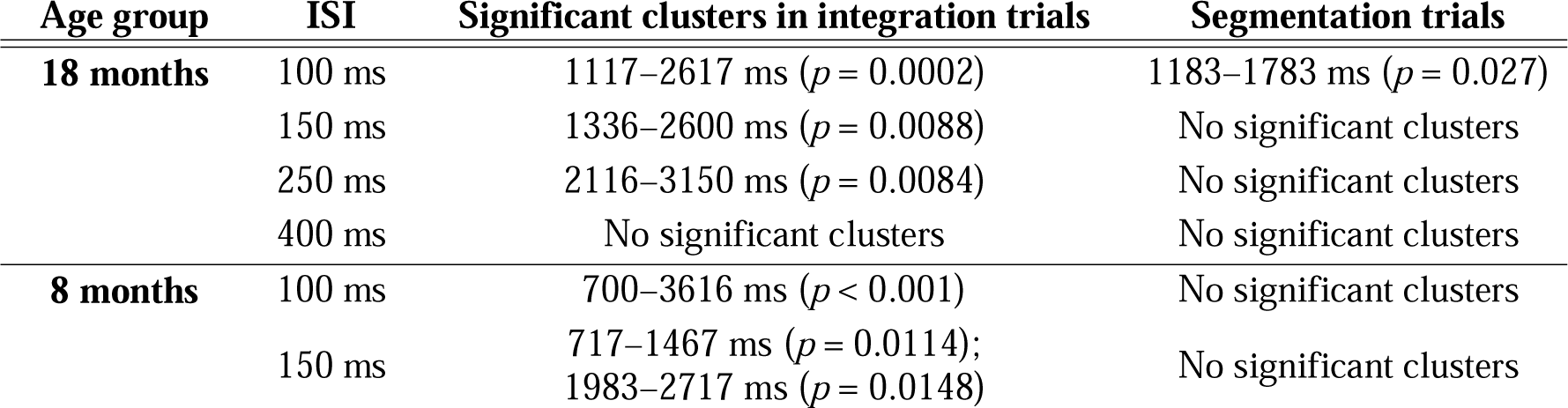
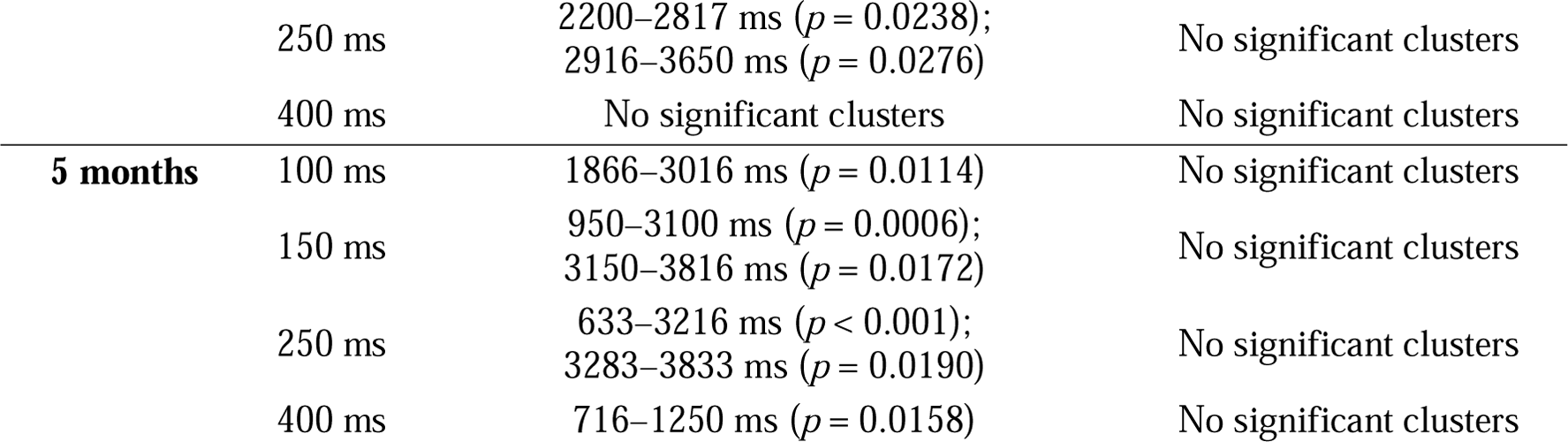
Evidence for target detection in Experiment 4.

## DISCUSSION

### Summary of the results

Using a novel instruction-free eye-tracking paradigm, we estimated the duration of the visual Temporal Integration Window (TIW) across six age groups from 5 months to adulthood. Across four experiments and 308 participants, we demonstrated a progressive and systematic shortening of the TIW over development (Figure 3), with estimates of approximately 400 ms at 5 months, ∼250 ms at 8 and 18 months, ∼100 ms at 3 years, and adult-like values of ∼57 ms by 5 years of age, a nearly sevenfold reduction. The steepest change occurred within the first year of life.

**Figure 3.**
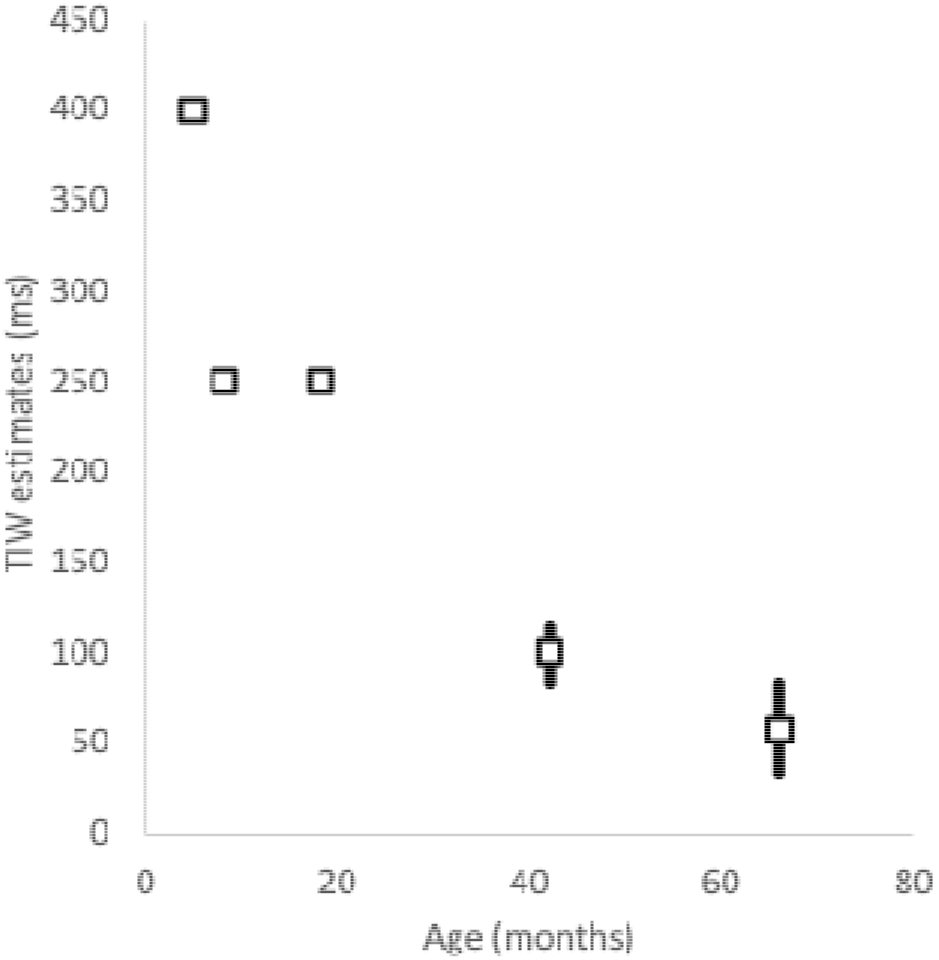
Estimation of the Temporal Integration Window (TIW) in ms, as a function of Age in months. Squares indicate the estimated TIW from infants’ and preschool children’s performance in integration trials. For 3- and 5-year-olds, black bars indicate the estimated range of TIW. The lower bound corresponds to the longest tested ISI for which the target was still reliably detected in integration trials, providing a minimum estimate of the TIW; the upper bound corresponds to the shortest tested ISI for which target detection failed, providing a maximum estimate of the TIW. For 8 and 18-month-olds the estimates constitute the upper bound. For 5-month-olds, the estimate constitutes the lower bound.

Our adult estimate of the TIW (∼57 ms) aligns with prior reports using variants of the same paradigm ^9,11^. Temporal integration with adults has also been studied across other sensory modalities and paradigms: using temporal order judgment tasks, Pöppel ^18,19^ reported an elementary processing threshold of approximately 30–40 ms that is invariant across vision, audition, and touch, while multisensory temporal binding windows for audiovisual pairs range from approximately 30–80 ms for simple stimuli to substantially wider windows for complex or speech stimuli ^20,21^. Our 5-year-old group showed adult-like TIWs in agreement with Freschl et al ^9^. Considering younger populations however, earlier work reported TIWs of about 2000 to 1000 ms in 6- to 15-month-old infants ^7^, substantially higher than our estimates for the same age range. This discrepancy likely reflects methodological differences across studies. But another interpretation is possible: even though we conceived integration and segmentation as two complementary processes, they may in fact constitute two independent processes. Notably, while our paradigm yielded informative data in integration trials at all ages, infants below 18 months did not detect the target in segmentation trials, even at ISIs that are longer than the estimated TIW (e.g., 400 ms at 18 months). Thus, there may be a temporal interval during which there is no more integration, but not yet efficient segmentation. Future studies could adjudicate this possibility, testing segmentation trials with longer ISIs.

Experiment 1 further demonstrated that the paradigm taps the pre-attentive stage of visual perception, as adults detected the target via a pop-out mechanism that was independent of the number of distractors, ruling out serial search ^22,23^.

Most importantly, we used the same paradigm to provide the first comprehensive and unified developmental trajectory of the visual TIW from early infancy to adulthood. More than the absolute values of the TIW at different developmental ages, we aimed at gaining information about the relative duration of the TIW across development. We observed a shrinking by about half in only 3 months of life, between 5 and 8 months, and by 7 until adulthood.

### Neural correlates of TIW maturation

The progressive shortening of the TIW aligns with the well-documented trajectory of brain structural maturation. The TIW is thought to reflect the efficiency of neural communication between visual processing areas, which is directly supported by white matter myelination and synaptic pruning ^11,24^. Myelination in the visual pathways increases steeply during the first postnatal months, then more gradually through childhood, with a direct impact on neural conduction speed ^25^. This structural trajectory is compatible with the pattern we observed: large TIW reductions in infancy, more moderate changes in childhood, and adult-like values by age 5. At the oscillatory level, peak alpha frequency, which tracks TIW duration in adults ^26–28^, gradually increases with age to reach adult values (∼10 Hz) around 4 years ^9,29^. This developmental increase in alpha frequency predicts a corresponding narrowing of TIWs, consistent with our finding that 5-year-olds already show adult-like temporal resolution. Although intrinsic neural timescales are distinct from behaviorally measured TIWs, longitudinal EEG findings reveal a parallel developmental pattern. These timescales shortened between 6 and 16 months and remained longer than in adults ^30^. The convergence between our behavioral trajectory and these neural markers strengthens the interpretation that TIW maturation reflects a core property of the developing visual system.

This developmental acceleration is not limited to the pre-attentive stage. The attentional blink, a measure of the temporal cost of the focused attention stage of visual processing, during which the system fails to process a second target following a first ^31^, reveals a strikingly similar developmental trajectory: from approximately 1150 ms at 5 months to ∼500 ms at 8 months and ∼200 ms at 3 years ^32,33^. The parallel shortening of both the TIW and the attentional blink suggests a broad and coordinated acceleration across the visual processing hierarchy, with the most pronounced developmental gains occurring between 5 and 8 months of age. This is further supported by fMRI evidence showing that neuronal intrinsic timescales are longer in neonates than in adults ^34^, and that 3- to 12-month-old infants segment the visual world into fewer, longer events than adults ^35^.

### Functional role of longer TIWs in early development

The extended TIWs observed in infancy may have important functional consequences for the content of early visual experience. A longer TIW means that more sensory information is integrated, possibly averaged, within each perceptual moment: transient and rapidly changing events are attenuated, while slow, sustained features of the environment are preserved. This low-pass temporal filtering may constitute an adaptive mechanism early in development, allowing infants to extract the stable, long-lasting structure of the world before the visual system is mature enough to resolve rapid changes. This interpretation is consistent with a broader framework in which the immature visual system acts as a set of natural perceptual filters, including low visual acuity and reduced color sensitivity, that constrain and sequence the information available to the infant, thereby supporting efficient perceptual learning ^36,37^. Long TIWs can thus be understood as one component of this broader developmental filtering system. Conversely, the progressive shortening of TIWs through development increases temporal resolution, enabling older children and adults to detect rapid changes and segment events at finer timescales, a capacity that is critical for processing dynamic social interactions, speech, and causal sequences ^38^.

## Conclusion

Across four experiments with same paradigm, we estimated that the temporal integration window at different developmental stages. We observed that the TIW decreases nearly sevenfold from early infancy to adulthood, shrinking from approximately 400 ms at 5 months of age to 57 ms in 5-year-olds and adults. These results point to a marked acceleration of visual processing over development, with the most dramatic changes occurring within the first year of life. Although the broader implications of this acceleration are only beginning to be understood, the present findings suggest that infants and adults perceive the world on fundamentally different temporal scales, differences that may give rise to profoundly distinct perceptual experiences.

## Acknowledgement

The current work was supported by Centre Hospitalier Le Vinatier, the ANR Grant ANR-25-CE28-5507-01 (JRH, MMGC), by the Fondation de France, grant number WB-2022-46104 (JRH), by LABEX CORTEX ANR-11-LABX-0042 of Université de Lyon (JRH).

## METHOD

### EXPERIMENTAL MODEL AND SUBJECT DETAILS

#### Experiment 1

Twenty-six neurotypical adults participated in the study (N = 26; 16 females; mean age = 27.16 years, SD = 5.68). Participants were recruited via flyers distributed in various settings, including universities, research center, sports clubs, and other public venues. The study was approved by the ethics committee (Comité de Protection des Personnes Sud-Ouest et Outre-Mer IV). All participants provided written informed consent prior to participation and were informed that they could withdraw at any time without consequence. The sample size was determined based on an a priori power analysis conducted in G*Power 3.1 ^39^ for a repeated-measures ANOVA (within-subject design). The analysis assumed a moderate effect size of *f* = 0.25, consistent with previous studies on temporal integration ^9^, an α=.05, desired power (1–β) = .95, an average correlation among repeated measures of 0.50. This analysis indicated that a total of N = 23 participants would be sufficient to detect effects of this magnitude. We therefore recruited 26 adults. All participants contributed at least one valid trial per condition and were thus included in the analysis.

#### Experiment 2

Fifty-five typically developing children were recruited and tested: 26 five-year-olds (14 females; mean age = 64.94 months, SD = 3.4 months) and 29 three-year-olds (13 females; mean age = 41.78 months, SD = 3.69 months). Three children in the three-year-old group were excluded from analyses because they did not contribute at least one valid trial per condition. The final sample included 26 five-year-olds and 26 three-year-olds. All participants were recruited through the consultation of birth records at the local city halls. This study was approved by the ethical committee ‘‘Comité de Protection des Personnes Sud Est II’’. Parents gave an informed consent before participating in the study and were informed that they could withdraw their consent at any time without any consequence. The sample size was decided following the same method as experiment 1.

#### Experiment 3

Ninety-one typically developing infants were recruited and tested: 27 eighteen-month-olds (15 females; mean age = 18.15 months, SD = 0.28 months), 30 eight-month-olds (10 females; mean age = 8.35 months, SD = 0.23 months), and 34 five-month-olds (14 females; mean age = 5.25 months, SD = 0.29 months). Four infants in the eighteen-month-old group, three in the eight-month-old group, and three in the five-month-old group were excluded from analyses because they did not contribute at least one valid trial per condition. The final sample included 23 eighteen-month-olds, 27 eight-month-olds, and 31 five-month-olds (N = 81). All participants were recruited through the consultation of birth records at the local city halls. This study was approved by the ethical committee ‘‘Comité de Protection des Personnes Sud Est II’’. Parents gave an informed consent before participating in the study and were informed that they could withdraw their consent at any time without any consequence. The sample size was decided following the same method as experiment 1.

#### Experiment 4

One hundred and sixty-four typically developing infants were recruited and tested: 57 eighteen-month-olds (30 females; mean age = 18.6 months, SD = 0.31 months), 53 eight-month-olds (27 females; mean age = 8.5 months, SD = 0.32 months), and 54 five-month-olds (20 females; mean age = 5.3 months, SD = 0.30 months). Nine infants in the eighteen-month-old group, three in the eight-month-old group, and three in the five-month-old group were excluded from analyses because they did not contribute at least one valid trial per condition. The final sample included 48 eighteen-month-olds, 50 eight-month-olds, and 51 five-month-olds (N = 149). All participants were recruited through the consultation of birth records at the local city halls. This study was approved by the ethical committee ‘‘Comité de Protection des Personnes Sud Est II’’. Parents gave an informed consent before participating in the study and were informed that they could withdraw their consent at any time without any consequence. Experiment 4 was designed to refine the infant TIW estimates obtained in Experiment 3 by testing longer ISIs. Infants contributed relatively few valid trials per condition. We therefore aimed to approximately double the sample size per age group relative to Experiment 3. This larger sample was intended to improve estimate precision and statistical sensitivity.

## METHOD DETAILS

### Stimuli

The stimuli were generated using MATLAB R2019b (MathWorks, Natick, MA) and displayed on a Tobii T60XL eye tracker screen with a resolution of 1920 × 1200 pixels. The stimuli consisted of 756 x 756 displays with grayscale faces or half-faces images (72 × 56 pixels) presented on a black background, and controlled for contrast and luminance. These grayscale faces were arranged along an invisible circle centred on the center of the screen, with a radius of 250 pixels. Two display disposition were used in Experiment 1: display with 12 faces (1 target among 11 distractors), with each face spaced every 30° along the circle; and display with 6 faces (1 target among 5 distractors) spaced every 60° (one item every two positions). In experiment 1, the position of the target could appear in all positions. For experiment 2, 3, and 4 only one display disposition was used (1 target among 11 distractors) and the position of the target could only appear in two positions, symmetrically located on the central axis of the screen (horizontal left and horizontal right).

At the nominal viewing distance of 60 cm, the 756 × 756-pixel stimulus field subtended approximately 19.3° × 19.3° of visual angle. The center of each face or half-face was located at approximately 6.4° eccentricity relative to the central fixation point. Because each trial began after central fixation was detected, the stimuli were initially presented outside the central fovea, within the near-peripheral visual field.

Pairs of displays A and B were created for integration trials and for segmentation trials. In integration trials, both Display A and Display B contained only half-faces. Display A contained half-faces in half of the positions (i.e., 6 or 3 positions). Display B contained half-faces in the positions not filled by the corresponding Display A, plus the complement of one half-face from corresponding Display A in the target location. The integration of Display A and Display B would permit the perception of one full face among half-faces; the segmentation of Display A and Display B would yield scattered half-faces. In segmentation trials, Display A contained half-faces at all non-target locations and no item at the target location. Display B contained the complementary half-faces at the non-target locations and a full face at the target location. The integration of Display A and Display B would yield the perception of full-faces all along the circle; the segmentation of Display A and Display B would yield the repeated appearance/disappearance of one full face among alternating half-faces (Figure 1.a-b, Figure S1).

### Procedure

Participants were seated 60 cm away from a Tobii T60XL eye tracker in a quiet, dark, soundproof room, with no other light source than the eye tracker screen. For child participants, the caregiver sat behind or beside the child. For infant participants, they sat on their caregiver lap. The caregiver was instructed to remain silent and avoid interacting with the child or infant during the experiment. Calibration of the eye-tracking system was conducted before the experiment began. Calibration points appeared sequentially in the four corners and center of the screen. Once the calibration was achieved, the experimenter manually initiated the experimental session.

Each trial began with a fixation cross displayed at the center of the screen. The trial started only when the participant successfully fixated this cross. Trials then consisted of a 4-second sequence of repeatedly alternating visual displays (Display A – black screen – Display B – black screen). The duration of the black screen separating A and B displays defined the interstimulus interval (ISI), which varied across four timing values: 16 ms, 32 ms, 83 ms, 116 ms (for experiment 1, 2 and 3) and 100 ms, 150 ms, 250 ms, 400 ms (for experiment 4). No instruction was given to participants other than watching the screen. The experiment consisted of 160 trials in total, divided among 8 experimental conditions (2 trial types × 4 ISIs), 20 trials each. Conditions were intermixed in a randomized order. Adult participants were expected to complete the full set of trials. For children and infants, the testing session ended when they no longer attended to the screen, even if the full set of trials had not been completed.

## DATA PROCESSING AND STATISTICAL ANALYSIS

### Trial Selection and Validity Criteria

Raw eye-tracking data were pre-processed to identify valid trials. For each trial, we extracted x and y gaze coordinates and pupil size over time. For all experiments, a trial was considered valid if the participant’s gaze remained within the screen area for at least 50 % of the samples during the first half of the trial (the first 2 seconds). This criterion ensured that analyses were based on trials where participants were effectively attending to the stimuli. All subsequent analyses were conducted exclusively on these valid trials.

### Participant Inclusion

Participants were retained in the final dataset only if they contributed at least one valid trial in each of the 8 experimental conditions (2 trial types × 4 ISIs). No participant in experiment 1 were excluded from the analysis. In experiment 2, 3 children of 3-year-old were excluded. In experiment 3, 4 infants in the 18 months old group, 3 infants in the 8 months old group and 3 infants in the 5 months old group did not meet this criterion and were excluded from the analysis. In experiment 4, 9 infants in the 18 months old group, 3 infants in the 8 months old group and 3 infants in the 5 months old group were excluded from the analysis.

### Interpolation of Missing Data

Gaze data from valid trials were processed using a custom correction function to handle missing samples. When missing x and y values occurred at the beginning or end of a trial, they were replaced by extrapolating the nearest available valid sample. Missing segments within a trial were filled using linear interpolation between adjacent valid samples. This approach preserves the temporal structure of the signal while minimizing discontinuities caused by brief tracking losses.

*Computation of Proportions toward Target vs. Distractors:*

For each trial, we extracted x–y gaze coordinates and identified gaze position within predefined regions of interest (ROIs). The target ROI was defined as a rectangular window centered on the target position ±100 pixels. In ***experiment 1***, distractor ROIs of the same size were defined at the other 5 or 11 stimulus positions. For each participant, at each time point, we computed the **Differential Looking Time (DLT)**, defined as:

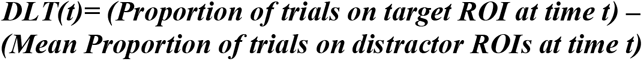

In ***experiment 2, 3 and 4*** ROIs were defined as a ±100-pixel window around the target and a symmetric distractor ROI on the opposite side. Again for each participant, at each time point, we computed the **DLT**, defined as

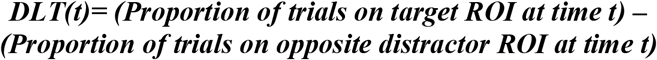

Positive values of DLTs indicate a tendency to look at the target. DLTs were computed for each condition and each participant, resulting in individual time courses. Statistical comparisons between conditions were performed using Cluster mass permutation test relying on ANOVAs and t-tests). To summarize participants’ performance, we calculated the mean DLT over the window of the meaningful effects of the trial for each participant and condition and conducted repeated measures ANOVAs to assess the effects of Trial Type (integration, segmentation) and ISIs (16, 32, 83, 116 ms in experiment 2 and 3; 100, 150, 250, 400 ms in experiment 4).

In experiment 1, to test whether the number of distractors affected target detection, we complemented the frequentist tests with Bayes factors. Bayes factors were computed for paired *t*-tests using the JZS default prior on effect size (Cauchy scale *r* = .707) ^40^. They are reported as BF10 when the data favored the presence of an effect and as BF01 when they favored its absence, following the conventional interpretation in which values between 1 and 3 are considered anecdotal, between 3 and 10 moderate, and above 10 strong.

## Supplementary materials

**Figure S1.**
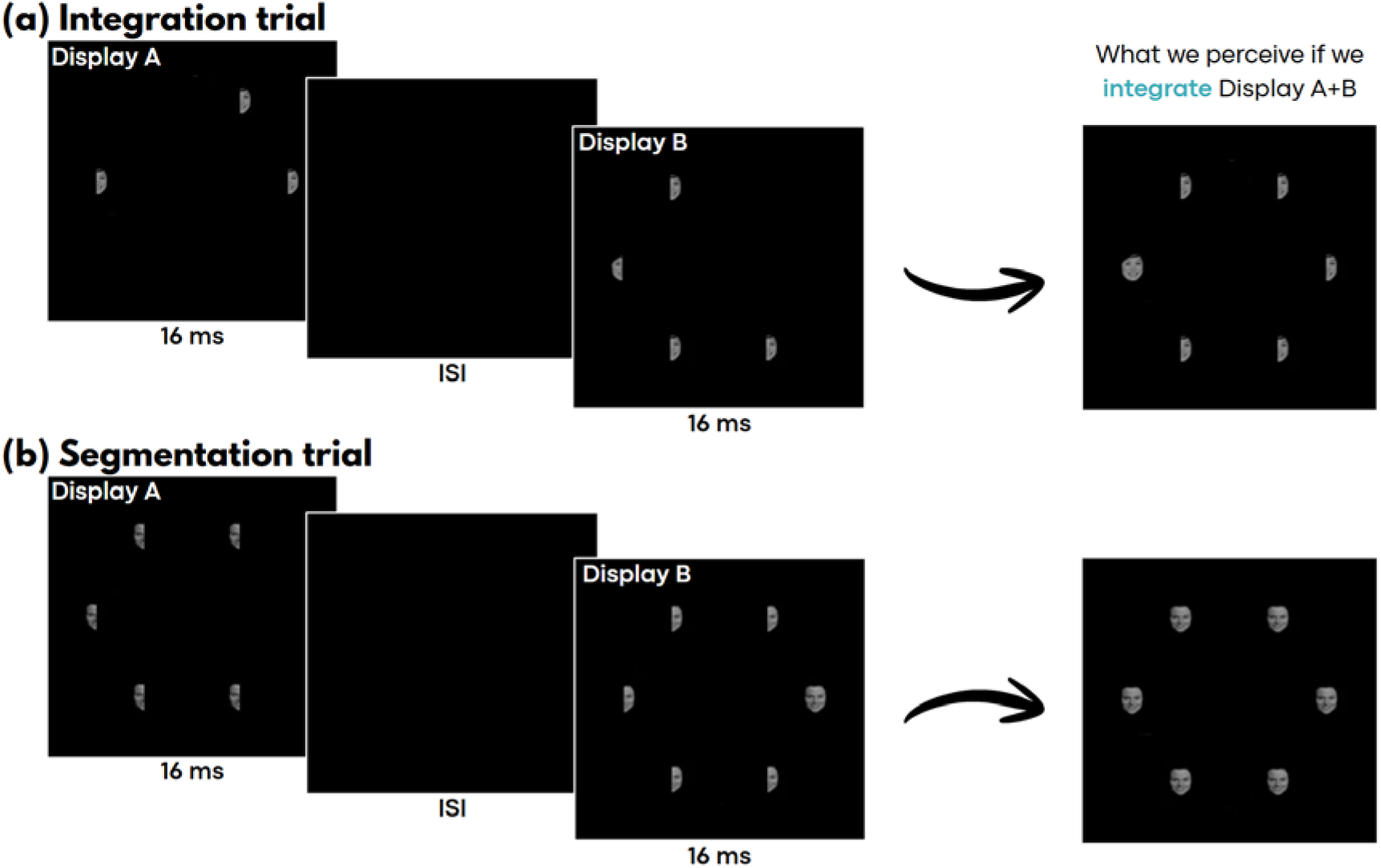
Paradigm Experiment 1 with 5 distractors **(a)** In integration trials, Display A and B alternate, presenting only half faces. Only two half faces complete each other, leading to the perception of a full face among half-faces when the two displays are integrated. **(b)** In segmentation trials, Display A contains half-faces at the non-target locations and no item at the target location, whereas Display B contains the complementary half-faces and one full face at the target location. Integrating the two displays produces full faces at all locations, whereas segmenting them reveals the single full face in Display B.

**Figure S2.**
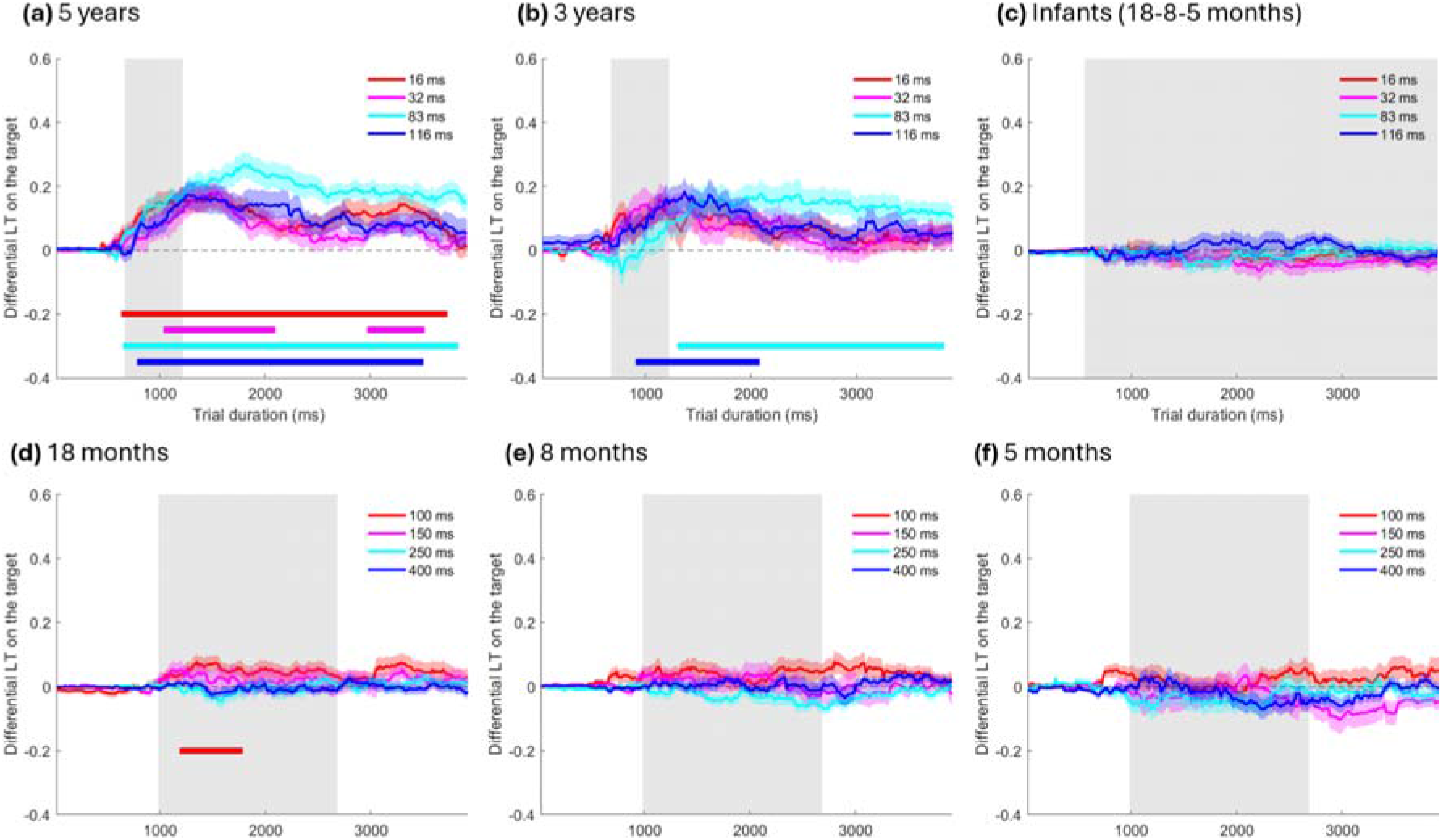
Differential Looking Times time course for segmentation trials in Experiments 2-4. **(a-c)** Results of Experiments 2 and 3. Target detection was assessed by comparing gaze toward the target location with gaze toward the opposite location. **(a)** Time course of target detection in segmentation trials for 5-year-olds (N = 26) at ISIs of 16, 32, 83, and 116 ms. **(b)** Time course for 3-year-olds (N = 26) at the same ISIs. **(c)** Time course for 5-, 8-, and 18-month-olds (total N = 81) at the same ISIs. In **(a)** and **(b)**, grey shading indicates the Age × Trial Type × ISI interaction window (666–1216 ms). In **(c)**, it indicates the interval showing a main effect of Trial Type. **(d-f)** Results of Experiment 4. **(d)** Time course of target detection in segmentation trials for 18-month-olds (N = 48), **(e)** 8-month-olds (N = 50), and **(f)** 5-month-olds (N = 51), at ISIs of 100, 150, 250, and 400 ms. In **(a-f)**, colored horizontal bars indicate significant time intervals for each ISI (cluster-mass permutation tests, p < 0.05). In **(d-f)**, grey shading indicates the window showing main effects of Trial Type and ISI (983–2683 ms).

---

**Table S1.** Mean number of trials presented and valid per condition across experiments and age groups.

| Condition | Number of trials presented | Number of valid trials |
| --- | --- | --- |
| **Adults (Experiment 1)** |  |  |
| Integration 16 ms | $19.92 \pm 0.08$ | $18.88 \pm 0.31$ |
| Integration 32 ms | $19.96 \pm 0.04$ | $18.96 \pm 0.29$ |
| Integration 83 ms | $19.81 \pm 0.12$ | $19.00 \pm 0.27$ |
| Integration 116 ms | $19.85 \pm 0.09$ | $18.96 \pm 0.32$ |
| Segmentation 16 ms | $19.92 \pm 0.05$ | $19.15 \pm 0.29$ |
| Segmentation 32 ms | $19.92 \pm 0.08$ | $19.00 \pm 0.28$ |
| Segmentation 83 ms | $19.85 \pm 0.07$ | $18.96 \pm 0.37$ |
| Segmentation 116 ms | $19.96 \pm 0.04$ | $19.12 \pm 0.32$ |
| **5-year-olds (Experiment 2)** |  |  |
| Integration 16 ms | $12.81 \pm 1.06$ | $11.73 \pm 1.00$ |
| Integration 32 ms | $12.85 \pm 1.11$ | $12.23 \pm 1.10$ |
| Integration 83 ms | $13.15 \pm 1.06$ | $12.00 \pm 0.99$ |
| Integration 116 ms | $12.77 \pm 1.15$ | $11.54 \pm 1.11$ |
| Segmentation 16 ms | $13.02 \pm 1.12$ | $12.19 \pm 1.04$ |
| Segmentation 32 ms | $12.62 \pm 1.15$ | $11.73 \pm 1.07$ |
| Segmentation 83 ms | $12.96 \pm 1.09$ | $11.96 \pm 1.05$ |
| Segmentation 116 ms | $12.38 \pm 1.05$ | $11.50 \pm 1.04$ |
| **3-year-olds (Experiment 2)** |  |  |
| Integration 16 ms | 11.31 ± 1.10 | 9.77 ± 0.96 |
| Integration 32 ms | 11.08 ± 1.11 | 9.69 ± 1.06 |
| Integration 83 ms | 11.19 ± 1.23 | 9.23 ± 1.11 |
| Integration 116 ms | 11.00 ± 1.15 | 9.35 ± 1.02 |
| Segmentation 16 ms | 11.46 ± 1.18 | 9.77 ± 1.11 |
| Segmentation 32 ms | 11.35 ± 1.08 | 9.27 ± 1.07 |
| Segmentation 83 ms | 11.35 ± 1.02 | 9.42 ± 0.93 |
| Segmentation 116 ms | 10.69 ± 1.13 | 8.92 ± 1.06 |
| **18-month-olds, fast presentation (Experiment 3)** |  |  |
| Integration 16 ms | 5.96 ± 0.68 | 5.52 ± 0.60 |
| Integration 32 ms | 5.61 ± 0.63 | 5.00 ± 0.58 |
| Integration 83 ms | 6.26 ± 0.83 | 5.96 ± 0.86 |
| Integration 116 ms | 5.83 ± 0.76 | 5.35 ± 0.77 |
| Segmentation 16 ms | 5.91 ± 0.77 | 5.57 ± 0.79 |
| Segmentation 32 ms | 5.87 ± 0.72 | 5.57 ± 0.74 |
| Segmentation 83 ms | 6.04 ± 0.68 | 5.61 ± 0.63 |
| Segmentation 116 ms | 5.78 ± 0.67 | 5.39 ± 0.67 |
| **8-month-olds, fast presentation (Experiment 3)** |  |  |
| Integration 16 ms | 6.63 ± 0.57 | 6.19 ± 0.58 |
| Integration 32 ms | 5.93 ± 0.59 | 5.48 ± 0.62 |
| Integration 83 ms | 6.33 ± 0.65 | 5.81 ± 0.62 |
| Integration 116 ms | 6.04 ± 0.53 | 5.59 ± 0.51 |
| Segmentation 16 ms | 6.19 ± 0.62 | 5.93 ± 0.62 |
| Segmentation 32 ms | 5.96 ± 0.53 | 5.48 ± 0.54 |
| Segmentation 83 ms | 5.96 ± 0.58 | 5.52 ± 0.52 |
| Segmentation 116 ms | 5.85 ± 0.59 | 5.59 ± 0.59 |
| **5-month-olds, fast presentation (Experiment 3)** |  |  |
| Integration 16 ms | 6.74 ± 0.45 | 5.97 ± 0.49 |
| Integration 32 ms | 5.45 ± 0.47 | 5.03 ± 0.49 |
| Integration 83 ms | 6.29 ± 0.52 | 5.58 ± 0.48 |
| Integration 116 ms | 5.65 ± 0.48 | 5.06 ± 0.45 |
| Segmentation 16 ms | 6.55 ± 0.54 | 6.13 ± 0.53 |
| Segmentation 32 ms | 6.10 ± 0.52 | 5.71 ± 0.50 |
| Segmentation 83 ms | 6.16 ± 0.48 | 5.72 ± 0.48 |
| Segmentation 116 ms | 5.48 ± 0.33 | 5.00 ± 0.35 |
| **18-month-olds, slow presentation (Experiment 4)** |  |  |
| Integration 100 ms | 7.38 ± 0.49 | 6.75 ± 0.49 |
| Integration 150 ms | 7.00 ± 0.50 | 6.42 ± 0.50 |
| Integration 250 ms | 7.21 ± 0.46 | 6.40 ± 0.48 |
| Integration 400 ms | 6.96 ± 0.40 | 6.06 ± 0.39 |
| Segmentation 100 ms | 7.23 ± 0.45 | 6.63 ± 0.44 |
| Segmentation 150 ms | 6.31 ± 0.44 | 5.77 ± 0.42 |
| Segmentation 250 ms | 6.29 ± 0.38 | 5.85 ± 0.39 |
| Segmentation 400 ms | 7.06 ± 0.38 | 6.50 ± 0.41 |
| **8-month-olds, slow presentation (Experiment 4)** |  |  |
| Integration 100 ms | 7.96 ± 0.52 | 7.36 ± 0.54 |
| Integration 150 ms | 7.56 ± 0.54 | 6.76 ± 0.52 |
| Integration 250 ms | 8.02 ± 0.49 | 7.42 ± 0.47 |
| Integration 400 ms | 8.06 ± 0.56 | 7.10 ± 0.53 |
| Segmentation 100 ms | 7.70 ± 0.51 | 7.00 ± 0.50 |
| Segmentation 150 ms | 7.42 ± 0.64 | 6.92 ± 0.60 |
| Segmentation 250 ms | 7.66 ± 0.57 | 6.96 ± 0.57 |
| Segmentation 400 ms | 7.98 ± 0.55 | 7.30 ± 0.55 |

| **5-month-olds, slow presentation (Experiment 4)** |  |  |
| --- | --- | --- |
| Integration 100 ms | 6.20 ± 0.44 | 5.67 ± 0.44 |
| Integration 150 ms | 5.84 ± 0.41 | 5.24 ± 0.41 |
| Integration 250 ms | 6.33 ± 0.46 | 5.57 ± 0.47 |
| Integration 400 ms | 6.33 ± 0.41 | 5.33 ± 0.39 |
| Segmentation 100 ms | 6.63 ± 0.44 | 5.96 ± 0.45 |
| Segmentation 150 ms | 5.61 ± 0.40 | 4.84 ± 0.39 |
| Segmentation 250 ms | 5.84 ± 0.37 | 5.20 ± 0.40 |
| Segmentation 400 ms | 6.49 ± 0.37 | 5.63 ± 0.38 |
| (mean ± SEM) |  |  |

**Table S2.** Number of participants who fixated the target at least once in all conditions.

|  |  |
| --- | --- |
| **Experiment 1:** |  |
| Adults | 26/26 |
| **Experiment 2: 3-5-year-olds** |  |
| 3-year-olds | 19/26 |
| 5-year-olds | 21/26 |
| **Experiment 3: 5-, 8-, and 18-month-olds, fast ISIs** |  |
| 18-month-olds | 8/23 |
| 8-month-olds | 15/27 |
| 5-month-olds | 13/31 |
| **Experiment 4: 5-, 8-, and 18-month-olds, slow ISIs** |  |
| 18-month-olds | 13/48 |
| 8-month-olds | 17/50 |
| 5-month-olds | 16/51 |

**Table S3.**
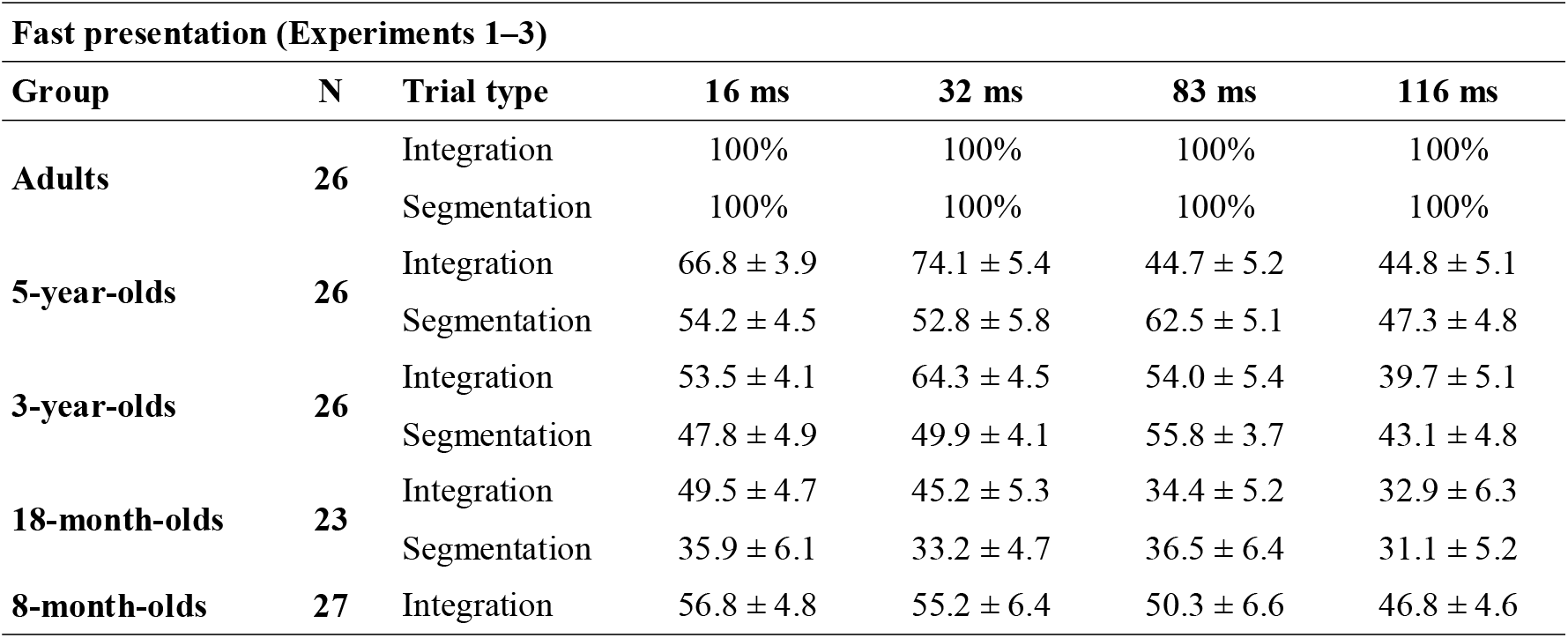
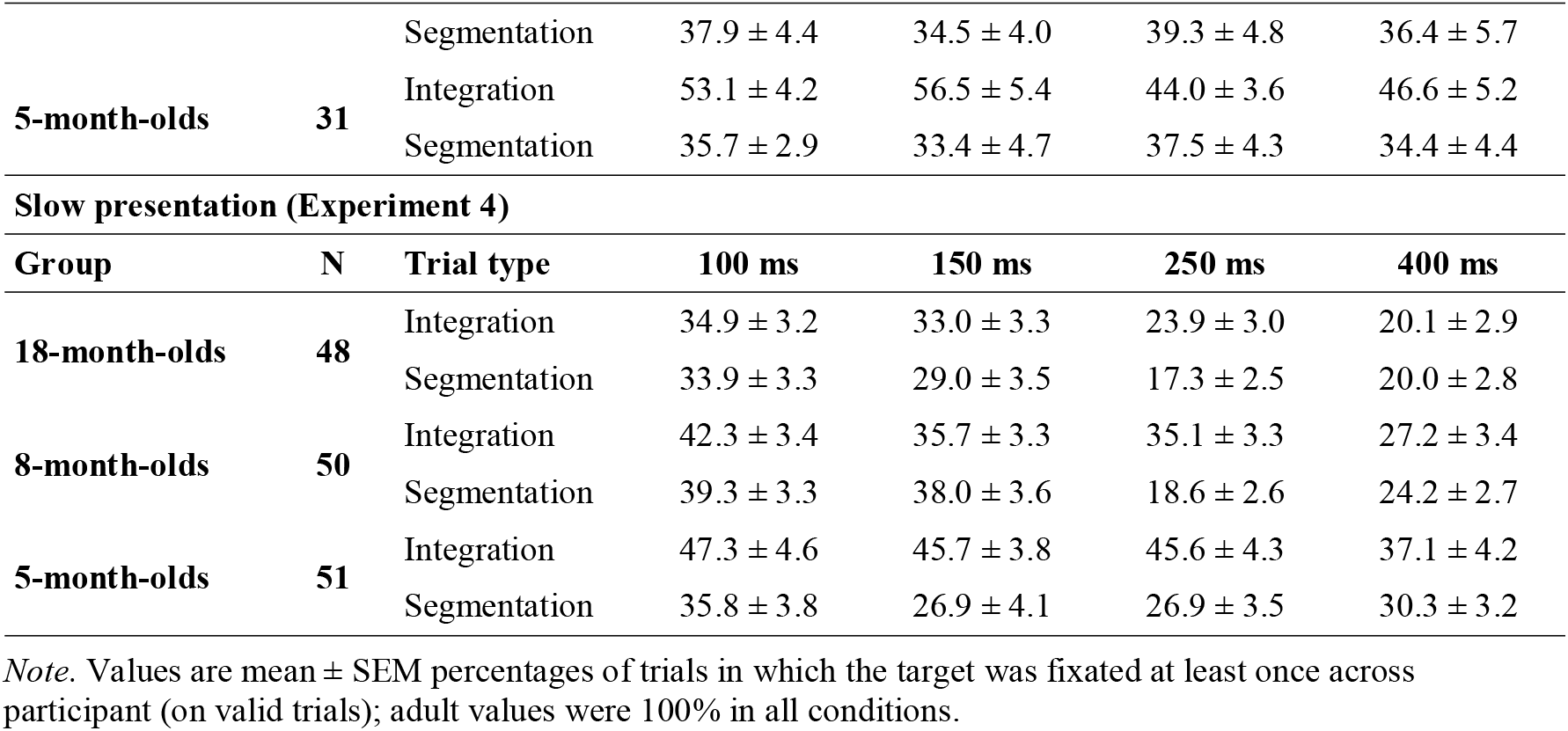
Mean percentage of valid trials in which the target was fixated at least once, by age group, trial type, and ISI.

## References

1. Stroud, J.M. (1949). The psychological moment in perception. In Transactions of the sixth conference on cybernetics, ed. Foerster, van. Macy, Josiah Jr.

2. Stroud, J.M. (1956). The fine structure of psychological time. In Information theory in psychology: problems and methods (Free Press), pp. 174–207.

3. Shallice, T. (1964). The Detection of Change and the Perceptual Moment Hypothesis. British Journal of Statistical Psychology 17, 113–135. 10.1111/j.2044-8317.1964.tb00254.x.

4. VanRullen, R. (2016). Perceptual Cycles. Trends in Cognitive Sciences 20, 723–735. 10.1016/j.tics.2016.07.006.

5. Goddu, M.K., and Gopnik, A. (2024). The development of human causal learning and reasoning. Nat Rev Psychol 3, 319–339. 10.1038/s44159-024-00300-5.

6. Michotte, A. (2017). The perception of causality (Routledge).

7. Farzin, F., Rivera, S.M., and Whitney, D. (2011). Time Crawls: The Temporal Resolution of Infants’ Visual Attention. Psychol Sci 22, 1004–1010. 10.1177/0956797611413291.

8. Freschl, J., Melcher, D., Kaldy, Z., and Blaser, E. (2020). Visual temporal integration windows are longer in infants. Journal of Vision 20, 1639. 10.1167/jov.20.11.1639.

9. Freschl, J., Melcher, D., Kaldy, Z., and Blaser, E. (2019). Visual temporal integration windows are adult-like in 5- to 7-year-old children. Journal of Vision 19, 5. 10.1167/19.7.5.

10. Freschl, J., Melcher, D., Carter, A., Kaldy, Z., and Blaser, E. (2021). Seeing a Page in a Flipbook: Shorter Visual Temporal Integration Windows in 2-Year-Old Toddlers with Autism Spectrum Disorder. Autism Research 14, 946–958. 10.1002/aur.2430.

11. Wutz, A., Muschter, E., van Koningsbruggen, M.G., Weisz, N., and Melcher, D. (2016). Temporal Integration Windows in Neural Processing and Perception Aligned to Saccadic Eye Movements. Current Biology 26, 1659–1668. 10.1016/j.cub.2016.04.070.

12. Davis, J., Hsieh, Y.-H., and Lee, H.-C. (2015). Humans perceive flicker artifacts at 500 Hz. Sci Rep 5, 7861. 10.1038/srep07861.

13. Treisman, A.M., and Gelade, G. (1980). A feature-integration theory of attention. Cognitive Psychology 12, 97–136. 10.1016/0010-0285(80)90005-5.

14. Brown, V., Huey, D., and Findlay, J.M. (1997). Face Detection in Peripheral Vision: Do Faces Pop Out? Perception 26, 1555–1570. 10.1068/p261555.

15. Hershler, O., and Hochstein, S. (2005). At first sight: A high-level pop out effect for faces. Vision Research 45, 1707–1724. 10.1016/j.visres.2004.12.021.

16. Nothdurft, H.-C. (1993). Faces and Facial Expressions do not Pop Out. Perception 22, 1287–1298. 10.1068/p221287.

17. VanRullen, R. (2006). On second glance: Still no high-level pop-out effect for faces. Vision Research 46, 3017–3027. 10.1016/j.visres.2005.07.009.

18. Pöppel, E. (1997). A hierarchical model of temporal perception. Trends in Cognitive Sciences 1, 56–61. 10.1016/S1364-6613(97)01008-5.

19. Pöppel, E. (2009). Pre-semantically defined temporal windows for cognitive processing. Philos Trans R Soc Lond B Biol Sci 364, 1887–1896. 10.1098/rstb.2009.0015.

20. Stevenson, R.A., Siemann, J.K., Schneider, B.C., Eberly, H.E., Woynaroski, T.G., Camarata, S.M., and Wallace, M.T. (2014). Multisensory Temporal Integration in Autism Spectrum Disorders. J. Neurosci. 34, 691–697. 10.1523/JNEUROSCI.3615-13.2014.

21. Vroomen, J., and Keetels, M. (2010). Perception of intersensory synchrony: A tutorial review. Attention, Perception, & Psychophysics 72, 871–884. 10.3758/APP.72.4.871.

22. Treisman, A.M., and Gelade, G. (1980). A feature-integration theory of attention. Cognitive Psychology 12, 97–136. 10.1016/0010-0285(80)90005-5.

23. Wolfe, J.M., O’Neill, P., and Bennett, S.C. (1998). Why are there eccentricity effects in visual search? Visual and attentional hypotheses. Perception & Psychophysics 60, 140–156. 10.3758/BF03211924.

24. Wutz, A., and Melcher, D. (2014). The temporal window of individuation limits visual capacity. Front. Psychol. 5. 10.3389/fpsyg.2014.00952.

25. Dubois, J., Dehaene-Lambertz, G., Perrin, M., Mangin, J.-F., Cointepas, Y., Duchesnay, E., Le Bihan, D., and Hertz-Pannier, L. (2008). Asynchrony of the early maturation of white matter bundles in healthy infants: Quantitative landmarks revealed noninvasively by diffusion tensor imaging. Human Brain Mapping 29, 14–27. 10.1002/hbm.20363.

26. Ronconi, L., and Melcher, D. (2017). The Role of Oscillatory Phase in Determining the Temporal Organization of Perception: Evidence from Sensory Entrainment. J. Neurosci. 37, 10636–10644. 10.1523/JNEUROSCI.1704-17.2017.

27. Samaha, J., and Postle, B.R. (2015). The Speed of Alpha-Band Oscillations Predicts the Temporal Resolution of Visual Perception. Curr Biol 25, 2985–2990. 10.1016/j.cub.2015.10.007.

28. Wutz, A., Melcher, D., and Samaha, J. (2018). Frequency modulation of neural oscillations according to visual task demands. Proceedings of the National Academy of Sciences 115, 1346–1351. 10.1073/pnas.1713318115.

29. Miskovic, V., Ma, X., Chou, C.-A., Fan, M., Owens, M., Sayama, H., and Gibb, B.E. (2015). Developmental changes in spontaneous electrocortical activity and network organization from early to late childhood. NeuroImage 118, 237–247. 10.1016/j.neuroimage.2015.06.013.

30. Truzzi, A., Rico-Picó, J., Rueda, M.R., and Cusack, R. (2026). The longitudinal development of intrinsic timescales in infancy and their relation to alpha brain rhythm. Cereb Cortex 36, bhag077. 10.1093/cercor/bhag077.

31. Raymond, J.E., Shapiro, K.L., and Arnell, K.M. (1992). Temporary suppression of visual processing in an RSVP task: An attentional blink? Journal of Experimental Psychology: Human Perception and Performance 18, 849–860. 10.1037/0096-1523.18.3.849.

32. Hochmann, J.-R., and Kouider, S. (2022). Acceleration of information processing en route to perceptual awareness in infancy. Current Biology 32, 1206–1210.e3. 10.1016/j.cub.2022.01.029.

33. Leroy, F., Naik, S., Gulbinaite, R., Palu, M., Battaglia, D., and Dehaene-Lambertz, G. (2026). Dynamics of the attentional blink in preverbal infants. Proceedings of the National Academy of Sciences 123, e2526752123. 10.1073/pnas.2526752123.

34. Truzzi, A., and Cusack, R. (2023). The development of intrinsic timescales: A comparison between the neonate and adult brain. NeuroImage 275, 120155. 10.1016/j.neuroimage.2023.120155.

35. Yates, T.S., Skalaban, L.J., Ellis, C.T., Bracher, A.J., Baldassano, C., and Turk-Browne, N.B. (2022). Neural event segmentation of continuous experience in human infants. Proc. Natl. Acad. Sci. U.S.A. 119, e2200257119. 10.1073/pnas.2200257119.

36. Jinsi, O., Henderson, M.M., and Tarr, M.J. (2023). Early experience with low-pass filtered images facilitates visual category learning in a neural network model. PLOS ONE 18, e0280145. 10.1371/journal.pone.0280145.

37. Spriet, C., Serraille, E., Naderi, S., Agrebi, M., Bouyahia, Z., Derrode, S., Kaiser, D., Papeo, L., and Hochmann, J.-R. Acceleration of vision changes object representations from infancy to adulthood. Preprint.

38. Kliesch, C., Parise, E., Reid, V., and Hoehl, S. (2022). The role of social signals in segmenting observed actions in 18-month-old children. Developmental Science 25, e13198. 10.1111/desc.13198.

39. Faul, F., Erdfelder, E., Lang, A.-G., and Buchner, A. (2007). G*Power 3: A flexible statistical power analysis program for the social, behavioral, and biomedical sciences. Behavior Research Methods 39, 175–191. 10.3758/BF03193146.

40. Rouder, J.N., Speckman, P.L., Sun, D., Morey, R.D., and Iverson, G. (2009). Bayesian t tests for accepting and rejecting the null hypothesis. Psychonomic Bulletin & Review 16, 225–237. 10.3758/PBR.16.2.225.

